# Discovery and in-vitro evaluation of potent SARS-CoV-2 entry inhibitors

**DOI:** 10.1101/2021.04.02.438204

**Authors:** Arpan Acharya, Kabita Pandey, Michellie Thurman, Elizabeth Klug, Jay Trivedi, Christian L. Lorson, Kamal Singh, Siddappa N. Byrareddy

## Abstract

SARS-CoV-2 infection initiates with the attachment of spike protein to the ACE2 receptor. While vaccines have been developed, no SARS-CoV-2 specific small molecule inhibitors have been approved. Herein, utilizing the crystal structure of the ACE2/Spike receptor binding domain (S-RBD) complex in computer-aided drug design (CADD) approach, we docked ∼8 million compounds within the pockets residing at S-RBD/ACE2 interface. Five best hits depending on the docking score, were selected and tested for their *in vitro* efficacy to block SARS-CoV-2 replication. Of these, two compounds (MU-UNMC-1 and MU-UNMC-2) blocked SARS-CoV-2 replication at sub-micromolar IC_50_ in human bronchial epithelial cells (UNCN1T) and Vero cells. Furthermore, MU-UNMC-2 was highly potent in blocking the virus entry by using pseudoviral particles expressing SARS-CoV-2 spike. Finally, we found that MU-UNMC-2 is highly synergistic with remdesivir (RDV), suggesting that minimal amounts are needed when used in combination with RDV, and has the potential to develop as a potential entry inhibitor for COVID-19.

## Introduction

Severe acute respiratory syndrome coronavirus 2 (SARS-CoV-2), the etiological agent of Coronavirus Disease 19 (COVID-19), emerged in early December 2019 in Wuhan City, China ^1^. Since its emergence, SARS-CoV-2 has claimed more than 2.6 million lives (https://coronavirus.jhu.edu/map.html) and caused unprecedented economic loss globally ^2^. Fourteen open reading frames (ORFs) of ∼ 30 kb positive (+) single-stranded SARS-CoV-2 RNA genome encodes up to sixteen non-structural proteins, four structural proteins, and nine accessory proteins ^3^. The spike glycoprotein, membrane protein (M), and E protein from the spherical envelop of SARS-CoV-2 ^4^. The spike protein (S protein) consists of S1 and S2 subunits ^5^. S1 subunit consists of an N-terminal domain (NTD) and a receptor-binding domain (RBD). In contrast, the S2 subunit has fusion peptide (FP), heptad repeat 1 (HR1), central helix (CH), heptad repeat 2 (HR2), connector domain (CD), transmembrane domain (TM), and a cytoplasmic tail (CT) ^6^. The RDB of the S1 subunit binds with the host cell receptor angiotensin-converting enzyme 2 (ACE2) to facilitate viral entry into the host cell ^7^, and the S2 subunit mediates membrane fusion ^8^. Although binding of spike RBD and ACE2 is a well-documented determinant of cellular entry, depending on cell types, the virions may enter the cell through clathrin-mediated endosomal pathways or clathrin-independent non-endosomal pathways ^9^.

The development of a vaccine against emerging and reemerging viruses is the most rational approach for containment of their rapid transmission ^10^. After the outbreak of SARS-CoV-2, several groups started vaccine development against the virus utilizing multiple vaccine development platforms such as mRNA based vaccines (Moderna, USA, and Pfizer-BioNTech, Germany), viral vector-based vaccines (University of Oxford-AstraZeneca; the UK and Johnson & Johnson; USA), recombinant protein-based vaccine (CoVaxx; USA), inactivated SARS-CoV-2 virus-based vaccine (Bharat Biotech; India and Sinovac Biotech; China), live attenuated SARS-CoV-2 based vaccine (Codagenix-USA and The Serum Institute of India-India collaboration) among others ^11^. The majority of them are in phase II/III trials, whereas a few have been granted emergency use authorization from USA-FDA ^12^. However, the newly emerging strains containing multiple mutations within the RBD may compromise the efficacy of these vaccines ^13^. Additionally, these vaccines effectiveness and durability remain unknown for the vulnerable population, including children, pregnant women, immunocompromised individuals, and people with comorbidities for COVID-19 ^14^.

Several therapeutic candidates have been repurposed against SARS-CoV-2 to control the rapid transmission and COVID-19 associated mortality and morbidity with mild to modest efficacy ^15^. Few of them now at different stages of clinical trials, including favipiravir (NCT04303299); ribavirin (NCT04392427), and sofosbuvir (IRCT20200128) to inhibit viral RNA-dependent RNA polymerase (RdRp), as well as drugs presumably acting on viral entry such as arbidol (NCT04260594) and hydroxychloroquine (NCT04355026), protease inhibitors lopinavir-ritonavir (NCT04276688) and darunavir (NCT04303299), IL-6 blockers tocilizumab (NCT04445272) and doxycycline (NCT04433078), JAK-1/2 inhibitors baricitinib (NCT04358614) and ruxolitinib (NCT04338958) among others ^16^. At present, only remdesivir (RDV: GS-5734) has been approved for the treatment of hospitalized COVID-19 patients ^17^. A randomized, double-blind, placebo-controlled clinical trial conducted by NIH found shorter hospital stay for patients with COVID-19 when treated with remdesivir than placebo controls (NCT04280705) ^18^. However, another multicenter, randomized clinical trial conducted by WHO reported no superior effect of the RDV on hospitalized COVID-19 patients (NCT04315948) ^19^. As per the current USA center for disease control (CDC) guideline, no specific therapy has been recommended for non-hospitalized and hospitalized mild to moderate COVID-19 patients who do not require oxygen support. For the hospitalized patients requiring supplemental oxygen (with or without ventilation), RDV alone or RDV with dexamethasone is recommended depending upon patient condition and availability of the drugs (www.covid19treatmentguidelines.nih.gov). These facts underscore the necessity of developing potent antiviral agents that may serve as prophylactic and therapeutic agents against COVID-19.

The development of an effective prophylactic against COVID-19 should preferentially target the viral S-RBD because the first step of the SARS-CoV-2 viral life cycle involves binding of S-RBD to the host ACE2 receptor. The crystal structure of S-RBD in complex with human ACE2 ^20^ has provided atomic details of the interaction interface ^20^. This complex’s analyses showed that the S-RBD/ACE2 interface contains pockets of suitable size that can target in silico drug discovery endeavors. We interrogated S-RBD/ACE2 interface in our computer-aided drug design (CADD) approach and docked ∼8 million compounds within the pockets residing at the S-RBD/ACE2 interface. The five best hits depending on the docking score and visual inspection were selected for testing their *in vitro* efficacy to block SARS-CoV-2 replication in cell-based assays. Two compounds (MU-UNMC-1 and MU-UNMC-2) blocked viral replication at sub-micromolar IC_50_ and are characterized in detail in a human bronchial epithelial cell line (UNCN1T) and Vero cells, including synergy with RDV.

## Results

### Screening library of compounds using computer-aided drug designing approach

The flow chart used for screening the library of compounds using a computer-aided drug design approach is depicted in **Fig. 1A**. Out of the 8 million compounds screened, we selected five compounds to test their efficacy in the cell-based assays. The IUPAC name, molecular structure, and glide score of the compounds are described in Table S1. The docked poses of the five compounds at the S-RBD/ACE2 interface are shown in **Fig. 1B-F**. MU-UNMC-1 and MU-UNMC-2 anchored with a Glide score of −9.8 and −11.6. Whereas the docking score of MU-UNMC-6, MU-UNMC-7, and MU-UNMC-8 were −7.6, −7.2, and −6.4, respectively. Compound MU-UNMC-1 was docked in a pocket formed by spike residues R403, E406, Y453, Y505, and ACE2 residues H34, E37 (**Fig. 1B**). Compound MU-UNMC-2 is docked in a pocket formed by spike residues R403, E405, E406, Y453, Y505, and ACE2 residues H34, E37, and D38 (**Fig. 1C**). Compounds MU-UNMC-6, MU-UNMC-7, and MU-UNMC-8 were also docked in the same pocket as MU-UNMC-1 (**Fig. 1D, 1E & 1F**). However, the significant difference in the docking poses of the compounds MU-UNMC-6, MU-UNMC-7, and MU-UNMC-8 and MU-UNMC-1 or MU-UNMC-2 was the exposure of the part of the molecules out of the S-RBD/ACE2 interface. It is clear from the binding of the two compounds that MU-UNMC-2 has more interactions than MU-UNMC-1.

**Figure 1.**
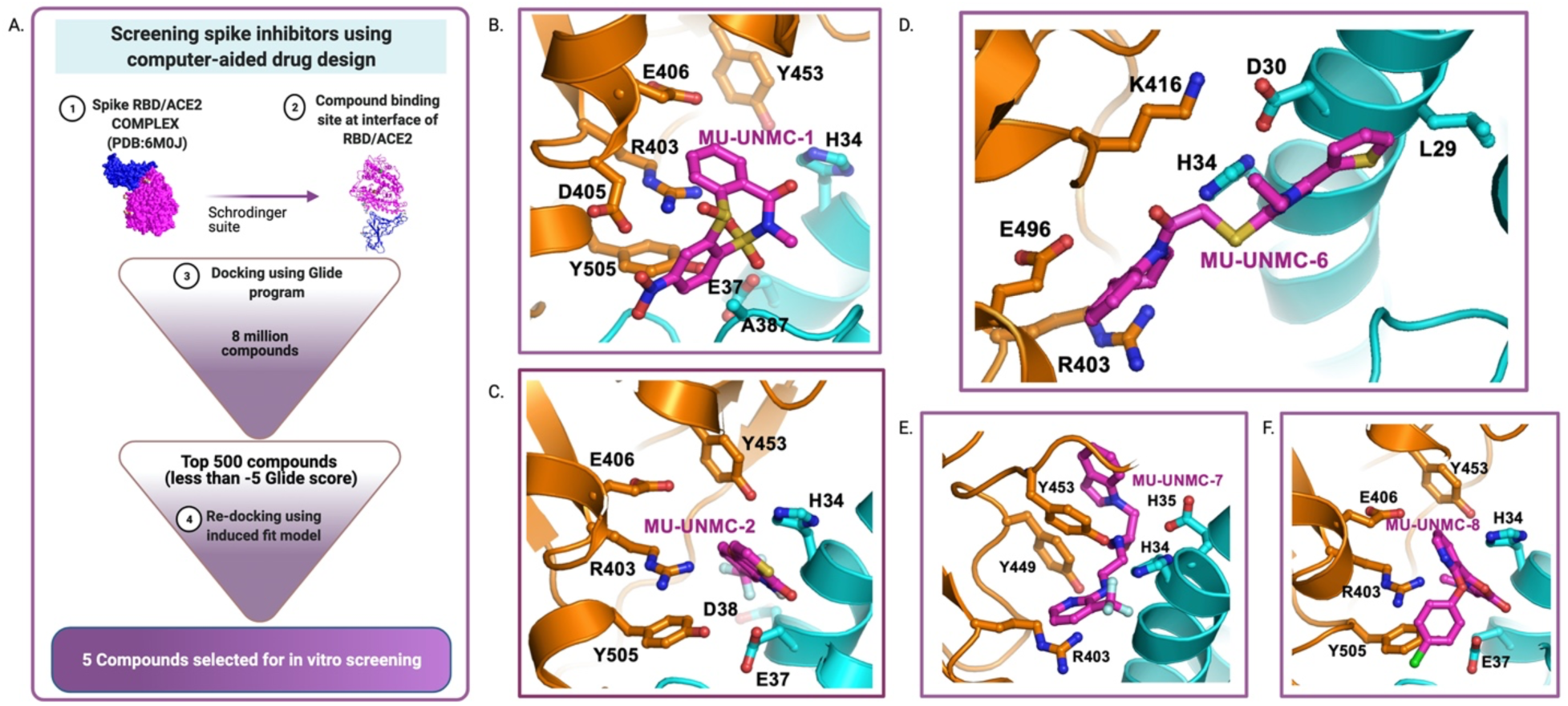
In-silico screening of SARS-CoV-2 entry inhibitors. (A) Flow chart of screening spike inhibitors using computer-aided drug designing approach. (B-F) Molecular docking of five compounds selected based on their glide score for in vitro screening of their antiviral activity against SARS-CoV-2.

### Determination of cellular toxicity of the compounds

The cytotoxicity of the five compounds was determined in Vero-STAT1 knockout cells using MTT assay described in the methods section. The percent viability of the cells is plotted against the increasing concentration of the compounds, and the fifty percent cytotoxic concentration (CC_50_) of each compound was computed using four-parameter variable slope sigmoidal dose-response models. The CC_50_ value of the compounds in Vero-STAT1 knockout cells was 6.21 μM for MU-UNMC-1 (**Fig. 2A**), 7.13 μM for MU-UNMC-2 (**Fig. 2B**), 37.18 μM for MU-UNMC-6 (**Fig. 2C**), 11.43 μM for MU-UNMC-7 (**Fig. 2D**) and 8.11 μM for MU-UNMC-8 (**Fig. 2E**) respectively.

**Figure 2.**
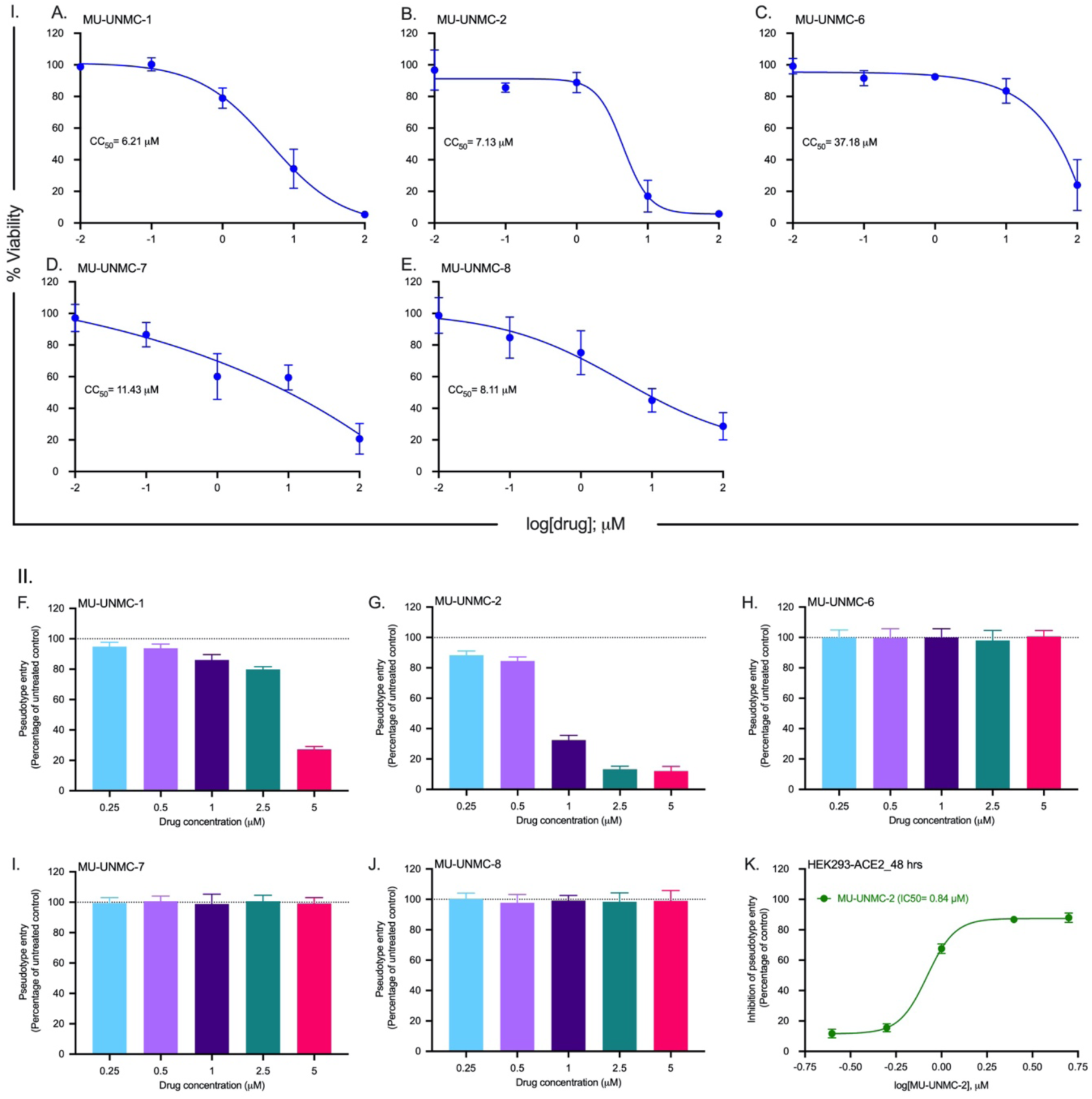
Measurement of cytotoxic concentration and screening of entry inhibition potential of five drug-like compounds using pseudovirus assay. Panel I: Measurement of 50% cytotoxic concentration (CC_50_) of five drug-like compounds. (A to E) Viability of Vero-STAT1 knockout cells in the presence of an indicated concentration of the compounds at 37°C for 72 hrs as measured by MTT assay. The CC_50_ values were computed using four-parameter variable slope sigmoidal dose-response models with Graph Pad Prism 8.0 software. Panel II: Screening entry inhibition potential of drug-like compounds in HEK293 cells expressing human ACE2. (F to J) HEK293-ACE2 cells were pretreated with the indicated concentration of compounds and then inoculated with pseudotyped lentiviral particles expressing spike glycoprotein of SARS-CoV-2. At 48 hrs post-inoculation, pseudotype entry was analyzed after normalization against untreated cells by determining luciferase activity in cell lysates. (K) The percentage inhibition of entry of pseudotyped lentiviral particles expressing spike glycoprotein of SARS-CoV-2 was measured for MU-UNMC-2 at indicated concentration of the compounds (cells that received DMSO were considered untreated controls). The IC_50_ value was computed using four parameter variable slope sigmoidal dose-response models using Graph Pad Prism 8.0 software.

### Screening entry inhibition efficacy of the compounds using lentiviral pseudovirus expressing SARS-CoV-2 spike protein

Replication of defective lentiviral particles expressing coronavirus spike glycoprotein effectively mimics coronavirus host cell entry ^21^. Therefore, we produced lentiviral particles representing the spike protein of SARS-CoV-2 on their surfaces as described in the Methods section and tested the entry blocking efficacy of the selected compounds using HEK293 cells engineered to express human ACE2. For this we first treated the HEK293-ACE2 cells with increasing concentrations of the compounds based on their CC_50_ values (0.25 to 5 μM), transduced them with pseudotyped lentiviral particles, and calculated the percentage of pseudotype entry against untreated controls (cells that received an equivalent amount of DMSO) after 48 hrs. We observed partially blocked pseudovirus entry by MU-UNMC-1 only at higher concentration (**Fig. 2F**) and robust inhibition pseudovirus entry by MU-UNMC-2 (**Fig. 2G).** However, the remaining three compounds MU-UNMC-6, MU-UNMC-7, and MU-UNMC-8, failed to prevent the pseudovirus entry **(Fig 2H, 2I & 2J).** Finally, we calculated the percentage inhibition of pseudotype entry by MU-UNMC-2 against untreated controls at increasing concentrations of the compound and the corresponding IC_50_ value (0.84 μM) using four-parameter variable slope sigmoidal dose-response models (**Fig. 2K**).

### Antiviral efficacy of MU-UNMC-1 and MU-UNMC-2 against wild type SARS-CoV-2

Next, we evaluated the antiviral efficacy of MU-UNMC-1 and MU-UNMC-2 against SARS-CoV-2 in a human bronchial epithelial cell line (UNCN1T), which is expected to demonstrate the in-vivo pathogenesis of SARS-CoV-2 in the lung and serve as a good model for the in-vitro efficacy study of antiviral compounds. We also use Vero-STAT1 knockout cells that are highly susceptible to viral infection due to the absence of cellular antiviral response.

The viral replication kinetics at 24 hrs and 48 hrs post-infection in the culture supernatant of SARS-CoV-2 infected UNCN1T, and Vero-STAT1 knockout cells in the presence of a different concentration of MU-UNMC-1 and MU-UNMC-2 is presented in **Fig. S1**. Based on SARS-CoV-2 viral loads in the culture supernatant of UNCN1T cells, MU-UNMC-1 and MU-UNMC-2 showed potent antiviral activity with an IC_50_ value of 0.67 μM and 1.72 μM at 24 hrs post-infection (**Fig. 3A**). In contrast, the compounds had an IC_50_ value of 1.16 μM and 0.89 μM respectively 48 hrs post-infection (**Fig. 3B**). On the other hand, in Vero-STAT1 knockout cells, 24 hrs post-infection MU-UNMC-1 and MU-UNMC-2 have an IC_50_ value of 5.35 μM and 1.63 μM (**Fig. 3C**), while at 48 hrs post-infection, the compounds have an IC_50_ value of 2.94 μM and 0.54 μM respectively (**Fig. 3D**).

**Figure 3.**
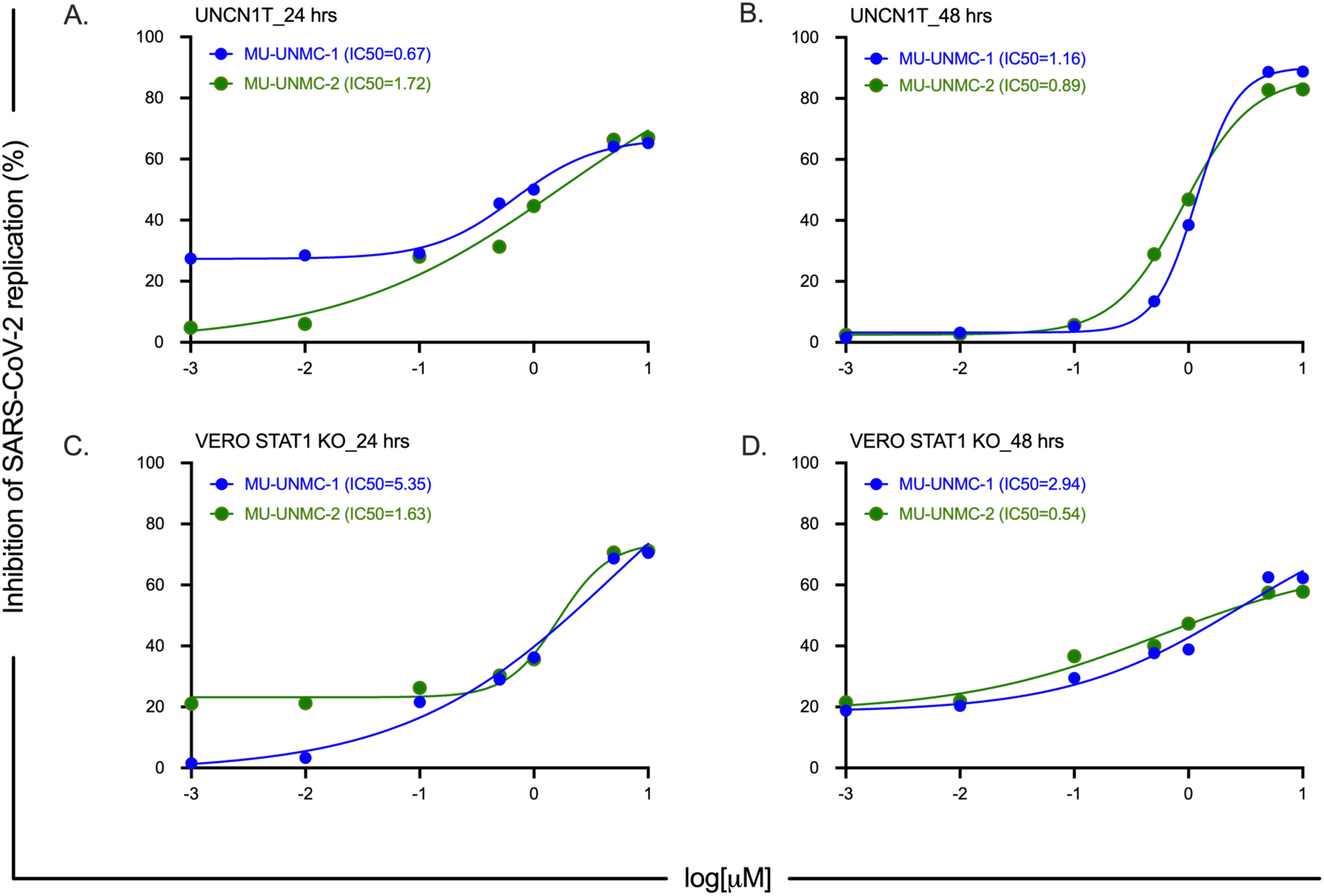
SARS-CoV-2 dose-response curve in MU-UNMC-1 and MU-UNMC-2 treated and SARS-CoV-2 infected UNCN1T and Vero-STAT1 knockout cells. (A & B) MU-UNMC-1 (in blue) and MU-UNMC-2 (in green) dose-response curve by percentage inhibition of SARS-CoV-2 replication 24 and 48 hrs post-infection in UNCN1T cells with indicated drug concentrations. (C & D) MU-UNMC-1 (in blue) and MU-UNMC-2 (in green) dose-response curve by percentage inhibition of SARS-CoV-2 replication 24 and 48 hrs post-infection in Vero-STAT1 knockout cells with indicated drug concentrations.

To understand the low activity of MU-UNMC-1 in blocking the entry of pseudovirus particles expressing SARS-CoV-2 spike glycoprotein while maintaining a potent antiviral activity against the wildtype virus, we revisited the molecular docking of this compound. We found that in-silico MU-UNMC-1 docked in two binding modes in the interface of spike and ACE-2 (**Fig. S2**). In the second binding mode, MU-UNMC-1 interacts with only a few residues. The two binding modes of the compound may be unstable at the interface of RBD and ACE2, leading to its low efficacy in pseudotyped assays.

### Chemical properties of MU-UNMC-2

We used the SwissADME web portal to compute the biophysical properties of all the compounds tested in this study. Here we discuss the properties of MU-UNMC-2, which showed the synergistic effect with RDV. The Log P_o/w_ values of this compound were 2.27 suggesting a high permeability of the compound. The predicted solubility of the compound was moderate. However, the pharmacokinetics data showed that the compound MU-UNMC-2 has high GI absorption. It was expected not to inhibit CYP2C9, CYP2D6, and CYP3A4 suggesting low toxicity of the compound. The compound followed all Lipnski’s Rules of five, and it was predicted as high drug likeliness with no PAINS (Pan-assay interference compounds).

### Combinational antiviral efficacy of MU-UNMC-2 and remdesivir

After finding robust effectiveness of MU-UNMC-2 in blocking the cellular entry of lentiviral-based pseudovirus particles expressing SARS-CoV-2 spike protein in HEK293-ACE2 cells and potent antiviral activity against wild type SARS-CoV-2 in UNCN1T cells, we evaluated the combinational antiviral effect of MU-UNMC-2 and remdesivir in UNCN1T and Vero-STAT1 knockout cells.

Based on mass action law and the classic model of enzyme-substrate inhibitor interactions, the median-effect equation was proposed by Chou (32), which describes the dose-effect relationship as (Fa/Fu) = (D/Dm)m or log(Fa/Fu) = mlog(D) – mlog(Dm). Where the slope (m) of the lines signifies the shape (m = 1, >1, and <1 signifies hyperbolic, sigmoidal, and flat sigmoidal dose-effect curves, respectively). Fa is defined as percentage inhibition of viral growth at dose D, and Fu is the fraction that remains unaffected (i.e., Fu = 1 - Fa). The antilog of x-intercept, where Fa/Fu = 1 or log(Fa/Fu) = 0, gives the Dm value. [log(Dm)] signifies the potency of the drugs (where Dm stands for median effective dose or IC_50_ concentration: the concentration required to inhibit 50% growth of the virus). Chou and Talalay proposed the combination index (CI) theorem to quantify synergism and antagonism between drug combinations ^22^. As per Chou and Talalay’s Combination Index Theorem, CI < 1, = 1, and >1 show synergism, additive effect, and antagonism, respectively. A plot of CI values at different fraction affected (Fa) levels is referred to as the Fa-CI plot or the Chou-Talalay plot.

Using CompuSyn, we generated the Median-Effect plot (Chou plot) of remdesivir (red line), MU-UNMC-2 (blue line), and the combination of doses of remdesivir and MU-UNMC-2 (green line) for SARS-CoV-2 infected UNCN1T cells 24 hrs post-infection as described above (**Fig. 4A**). Analysis of the Median-Effect, where the X-intercept of the lines indicates potency, shows that the combination dose is more potent than the individual doses of the compounds. In contrast, the Chou-Talalay plot (**Fig. 4B**) indicates the synergistic effect of remdesivir and MU-UNMC-2. The dose-response percent inhibition matrix of single and combined treatment of remdesivir and MU-UNMC-2 was computed using SynergyFinder v.2 (**Fig. 4C**). The 3-D interaction landscape between remdesivir and MU-UNMC-2 was computed based on Loewe additive model using SynergyFinder v.2 in SARS-CoV-2 infected UNCN1T cells 24 hrs post-infection (Loewe synergy score 26.64; with most synergistic area score of 37.25) (**Fig. 4D**). Synergy maps highlight synergistic and antagonistic dose regions in red and green colors, respectively.

**Figure 4.**
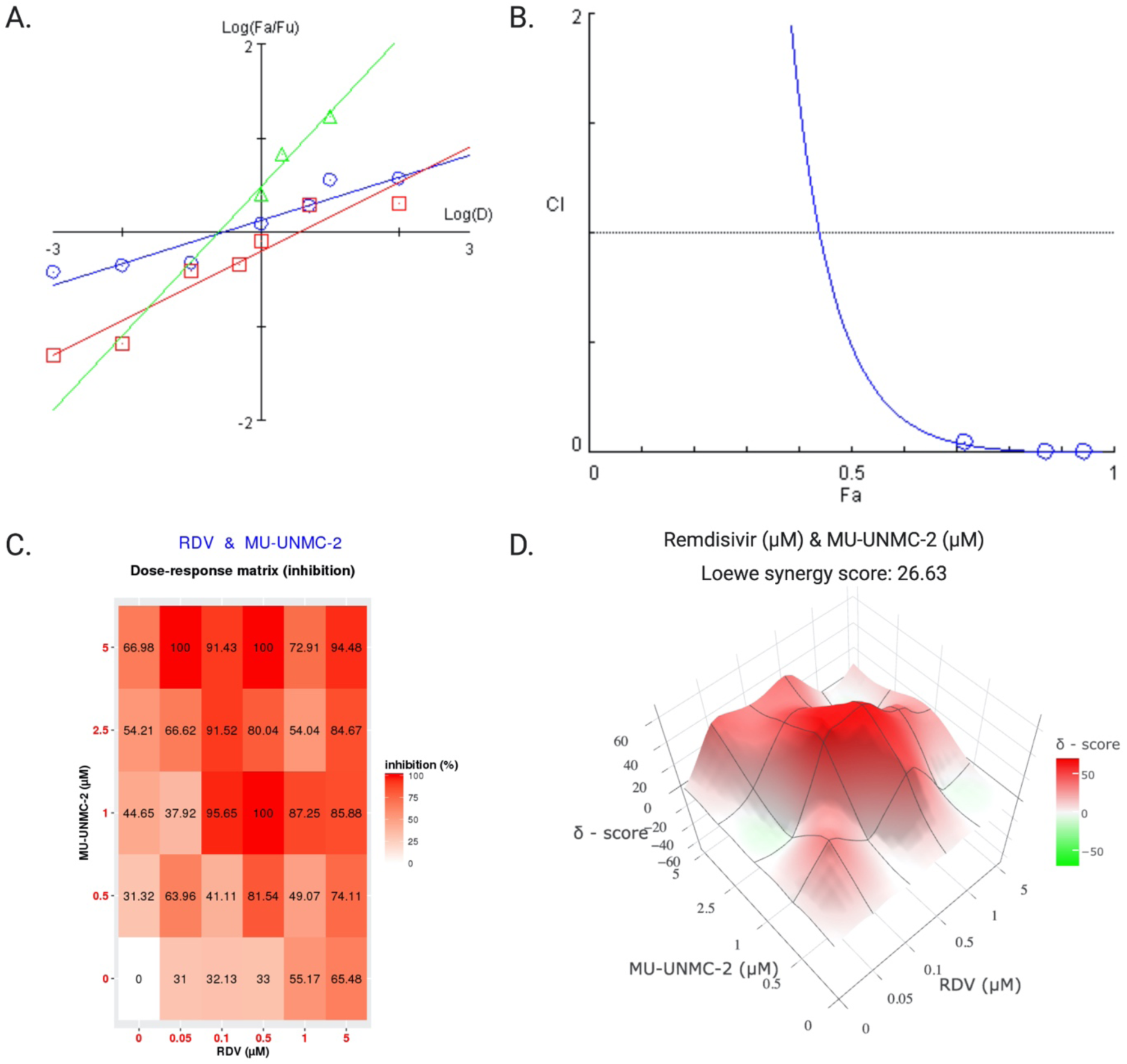
Synergistic effect of remdesivir and MU-UNMC-2 combined treatment against SARS-CoV-2 infected UNCN1T cells 24 hrs post infection. (A) The Median-Effect plot (Chou plot) of MU-UNMC-2 (the red line with square data points), remdesivir (blue line with circular data points) and fixed-dose combination of remdesivir and MU-UNMC-2 (green line with triangular data points). The median-Effect equation, that describes the dose-effect relationship is given by (Fa/Fu) = (D/Dm)^m^ or log(Fa/Fu) = m log(D) – m log(Dm). Fa is defined as percentage inhibition of viral growth at dose D and Fu is the fraction that remains unaffected (i.e., Fu = 1 - Fa). (B) Combination index plot or Chou-Talaly plot or Fa-CI plot: A plot of CI (combination index) on the Y-axis vs. Fa on X-axis for fixed dose combination of Remdesivir and MU-UNMC-2 in SARS-CoV-2 infected UNCN1T cells (blue line with circular data points). (C) Dose-response percent inhibition matrix of single and combined treatment of remdesivir and MU-UNMC-2 in SARS-CoV-2 infected UNCN1T cells 24 hrs post infection. (D) 3-D interaction landscape between remdesivir and MU-UNMC-2 calculated based on Loewe additive model using SynergyFinder v.2 in SARS-CoV-2 infected UNCN1T cells 24 hrs post-infection (Loewe synergy score 26.63; with most synergistic area score of 37.25).

Similarly, for SARS-CoV-2 infected Vero-STAT1 knockout cells treated with different combination doses of remdesivir and MU-UNMC-2, the Median-Effect plot (Chou plot) of remdesivir (blue line), MU-UNMC-2 (red line), and the fixed-dose combination of remdesivir and MU-UNMC-2 (green line) are described in **Fig. S3A**, the Chou-Talalay plot (**Fig. S3B**) indicates a borderline synergistic effect of remdesivir and MU-UNMC-2. The dose-response percent inhibition matrix of single and combined treatment of remdesivir and MU-UNMC-2 were computed using SynergyFinder v.2 (**Fig. S3C**). The 3-D interaction landscape between remdesivir and MU-UNMC-2 was derived using Loewe additive model and SynergyFinder v.2 in SARS-CoV-2 infected UNCN1T cells 24 hrs post-infection (Loewe synergy score 8.18; with most synergistic area score of 13.24) (**Fig. S3D**). **Table S2** shows the median effective dose of MU-UNMC-2, remdesivir, and their fixed-dose combinations. At the same time, **Table S3** shows the CI values at ED_50_, ED_75_, ED_90_, and ED_95_ in Vero-STAT1 knockout and UNCN1T cells, which suggests a minimal to solid synergistic effect of the drug combinations. In Vero-STAT1 knockout cells at Fa 0.5, we obtained a dose reduction index (DRI) of 3.84 for MU-UNMC-2 and 1.39 for remdesivir. On the other hand, in UNCN1T cells, we received a dose reduction index (DRI) of 28.30 for MU-UNMC-2 and 2.33 for remdesivir.

## Discussion

By utilizing *in silico* workflow, we screened an in-house library of the drug-like compounds containing ∼8 million compounds for their potential to bind at the interface of SARS-CoV-2 spike glycoprotein human ACE2 receptor. Based on their glide score and visual inspection, we selected five compounds to test their efficacy in SARS-CoV-2 replication cell-based assays. Out of the five, one compound (MU-UNMC-2) showed a potent entry inhibition in lentiviral-based pseudovirus assays. Whereas another compound (MU-UNMC-1) showed only a modest inhibition activity against virus entry. The other three compounds (MU-UNMC-6, MU-UNMC-7, and MU-UNMC-8) did not block either virus entry or replication. The antiviral activity of the two compounds active in pseudovirus assays was also tested to block viral replication in a physiologically relevant human bronchial epithelial cell line (UNCN1T). The 2-D *in vitro* culture system, MU-UNMC-1 and MU-UNMC-2, both show potent antiviral activity with IC_50_ values in the sub-micromolar range. These results showed that both MU-UNMC-1 and MU-UNMC-2 are SARS-CoV-2 inhibitors, but MU-UNMC-2 acts at the entry of the virus into the host cell and is found to be highly synergistic with remdesivir.

After the elucidation of the crystal structure of SARS-CoV-2 spike protein S-RBD bound to ACE2, several groups targeted this interface to develop entry inhibitors, including peptide analogs, monoclonal antibodies, and protein chimeras, among others ^23–31^ using *in silico* as well as *in vitro* settings. Despite the availabilities of these new technologies and macromolecules as therapeutic agents, small molecules have been a preferred choice for drug development for their better pharmacokinetic profile, stability, and dosage logistics ^32, 33^. Also, small molecules have the advantage of logistic chain shipment to remote areas, and their production cost is economical compared to peptide-based therapeutics. However, several studies are underway to develop small molecules and peptide inhibitors to block the human ACE2 ^34–37^. As ACE2 is a principal regulator of the renin-angiotensin system (RAS) that regulates the cardiovascular and renal system, ACE2 is not a preferable therapeutic target, as inhibition of this receptor may disrupt the physiological homeostasis of the systems mentioned above ^38^. Additionally, the ACE2/S-RBD interface is distal to the protease active site of ACE2. Hence, inhibitors developed against the ACE2 protease activity would most likely not affect S protein binding to ACE2. ACE2 plays a protective role in lung injury in acute respiratory distress syndrome (ARDS) caused by respiratory viral infections ^39^. In this respect, our strategy of screening small molecules that block S protein and ACE2 is a better approach and has an edge against protein-based therapeutics.

In the arena of antiviral therapeutics, the use of combination/repurposing drugs that target different stages of the viral life cycle and host factors is well documented, resulting in superior virological and physiological response compared to monotherapy ^40, 41^. This approach increases the overall efficacy of the treatment, reduces the dosages requirement of individual drugs, reduces their toxicity profile, and lowers the chances of developing drug resistance ^42^. This approach is time-tested during the development of combinational antiretroviral therapy (cART) to treat human immunodeficiency virus-1 (HIV-1) ^41, 43^. During the early days of the COVID-19 pandemic, investigational combination therapy approaches have been tested in severely ill hospitalized patients. These strategies only yielded mild to moderate responses in different clinical settings ^44, 45^. In a similar pursuit, we examined the combination of RDV, an antiviral known to inhibit viral RdRp, with MU-UNMC-2, which acts as an entry inhibitor in the *in vitro* assays using human bronchial epithelial cell line and wildtype SARS-CoV-2. Our finding showed a 28.3-fold and 2.3-fold reduction in dosages for MU-UNMC-2 and RDV, respectively, thereby supporting their combinatorial use of the two compounds against monotherapies. This indicates combination therapy targeting two critical stages of the viral life cycle, namely cell entry (MU-UNMC-2) and replication (RDV), enhances the overall therapeutic efficacy and reduces chances of development of resistance associated with one drug. The reduction in remdesivir dose is also expected to reduce its adverse side effects that substantially limit its use in clinical settings ^18, 46^.

As documented in the literature, the 2-D culture system is not the best model for screening antivirals as it does not reflect the *in vivo* physiological conditions. Nonetheless, it is an essential tool for initial screening prior to pursuing preclinical trials. Therefore, our future studies are focused on validating our findings in an appropriate animal model for COVID-19 ^47^.

Our strategy that combines *in silico* drug screening followed by *in vitro* testing in the cell-based system has a significant advantage over the classical drug-discovery approaches. It substantially reduces the resource and expenses for initial screening of drug-like compounds, increases the probability of selecting the best lead compounds with desired therapeutic profile with fewer trials. Our strategy also shortens the time between the lead identification-to optimization, preclinical test, to translational application to clinical trials. While we have not optimized the lead compound, we are in the process of doing so, and the results will be presented elsewhere. However, initial theoretical estimation of the ADME profile suggests that the compound MU-UNMC-2 has a favorable physicochemical profile and can be quickly taken into pre-clinical trials. Once the ADME profile is confirmed in suitable models, we expect that MU-UNMC-2 can be used in clinical trials either as monotherapy or combined with RDV. Since 122 million people have been infected and there have been 2.6 million deaths due to COVID-19, a prophylactic and therapeutic treatment option for the general population is urgently needed. With the emergence of new variants of the SARS-CoV-2 that are supposed to have enhanced/accelerated transmission, new potent antivirals are warranted. With the rolling out of few vaccines for EAU, there is the hope of light that the pandemic will be under control soon, though without a precise estimation of a timeline ^48^. Within this context, our present work is highly significant and can move to the clinic. The study’s result affirms the utility of computer-aided drug designing approaches in carefully conducted *silico* screening of an extensive library of compounds in preliminary screening for drug-like compounds for therapeutic use, which would otherwise take several years and billions of investments if tested by conventional drug screening techniques ^49^.

Herein we report *in silico* screening of drug-like compounds to find potential SARS-CoV-2 entry inhibitors. Out of the five best hit compounds, one molecule (MU-UNMC-2) alone or in combination with RDV showed potent antiviral activity against SARS-CoV-2 at micromolar concentrations. Additionally, this compound’s ADME toxicity properties were well within the criteria to fall in the category of drug-likeliness. Hence, MU-UNMC-2 may potentially be a prophylactic/therapeutic drug against COVID-19 once validated *in vivo*.

## Acknowledgments

We acknowledge the UNMC BSL-3 core facility for allowing us to perform all in vitro experiments involving SARS-CoV-2. ‘The University of Nebraska Medical Center BSL-3 Core Facility is administrated through the Office of the Vice Chancellor for Research and supported by the Nebraska Research Initiative (NRI).’ “The following reagent was deposited by the Centers for Disease Control and Prevention and obtained through BEI Resources, NIAID, NIH: (a) SARS-Related Coronavirus 2, Isolate USA-WI1/2020, NR-52384 and (b) Quantitative PCR (qPCR) Control RNA from Heat-Inactivated SARS-Related Coronavirus 2, Isolate USA-WA1/2020, NR 52347.”

## Author contributions

SNB conceive the study, data interpretation and edited the manuscript; KS design and performed the docking analysis and edited the manuscript; AA performed the experiments, analyze the data wrote the manuscript; JT performed the cytotoxicity analysis; MT, KP and EK performed viral infection and PCR analysis.

## Star Methods

### Identification of potential inhibitor using computer-aided drug design

To identify potential SARS-CoV-2 entry inhibitors, we used the docking of ∼8 million drug-like compounds from an in-house library of compounds. This library has been curated to include the compounds that follow Lipinski’s rule of five. The library consists of compounds from MayBridge Hitfinder, Zinc database ZincDatabase, Zinc15Database, ChEMBL, Bingo, JChemforExcel, ChemDiff, and BindingMOAD (https://www.click2drug.org/index.php#Databases). RBD/ACE2 complex (Protein Databank Entry 6M0J; Lan et al. 2020 ^20^) was used for *in silico* screening. A docking-ready structure was generated by ‘Protein Preparation Wizard’ of the Schrödinger Suite (Schrödinger LLC, NY), which adds the hydrogen atoms, missing sidechains, and assigns protonation states to His, Gln, and Asn together with the optimization of the hydrogen atoms’ orientation. The resulting structure was energy minimized using OPLS_2005 forcefield for 10,000 iterations to remove steric conflicts. Potential compound binding sites were identified by SiteMap (Schrödinger Suite). We selected the binding pockets present at the interface of S-RBD/ACE2 for the docking of the library compounds. The glide program of Schrödinger Suite with SP (Simple Precision) was first used for docking. The top 500 compounds, the best Glide, was then re-docked using the ‘Induced Fit’ program of the Schrödinger Suite. The resultant docking poses were visually inspected for their feasibility to inhibit the binding of RBD to ACE2. Five compounds were finally selected for testing their *in vitro* inhibitory activity in cell-based assays (**Table 1**).

### Chemical Properties of the compounds

In order to assess chemical properties of the compounds, we used the SwissADME web portal as detailed in Daina et al. ^50–52^. The input data were provided in the form of a SMILES notation. This web portal computes the biophysical properties, lipophilicity, water-solubility, pharmacokinetics, and drug-likeliness.

### Reagents and cell lines

Remdesivir (GS-5734) was obtained from Selleck Chemicals LLC(Houston, TX). The SARS-CoV-2 entry inhibitors were obtained from MolPort (Riga, Latvia). Vero E6 (CRL-1586™) and Vero-STAT1 knockout cells (CCL-81-VHG™) were obtained from ATCC®. These cells were cultured in DMEM containing 10% fetal bovine serum, 2 mM L-glutamine, 100 units/ml penicillin, 100 units/ml streptomycin, and 10 mM HEPES (pH 7.4). UNCN1T cells (a human bronchial epithelial cell line; Kerafast; cat# ENC011) were cultured in BEGM media (Bronchial Epithelial Cell Growth Medium; Lonza: cat# CC-3170) in FNC (Athena Enzyme Systems; cat# 0407) coated 96-well plates. All other reagents (molecular biology grad fine chemicals) used in the study were purchased from Sigma-Aldrich (St. Louis, MO) unless otherwise mentioned.

### MTT cell viability assay

Vero and HepG2 cells were seeded at the density of 20,000 cells/well in a 96 well plate containing 100 µL complete DMEM (Gibco, USA) supplemented with 10% FBS (Gibco, USA) and 1% Penstrep (Gibco, USA). Cells were incubated for 12 hours at 37° C in humidified 5% CO_2_ incubator for adherence. After 12-hour incubation, the media was replaced with fresh media, and cells were treated with the compound at concentrations ranging between 0.001 to 100 µM. Untreated cells were considered as negative control, and DMSO treated cells were considered as the vehicles. After the treatment, cells were incubated at 37 ° C in humidified 5% CO_2_ incubator. 48-hour post-treatment, 20 µL of MTT substrate (5 mg/mL) was added in each well and incubated for 4 additional hours at 37° C in the dark. 4 hpi, the media was carefully removed, and blue formazon crystals were dissolved in 200 µL of DMSO, and the purple color was read at 595 nm.

### Production and titration of lentiviral-based pseudoviruses expressing SARS-CoV-2 spike glycoprotein

For pseudotyping, lentiviral particles expressing SARS-CoV-2 spike protein were generated as described by Crawford et al. ^53^. In brief, 3X10^6^ HEK293T cells were co-transfected with a plasmid containing lentiviral backbone expressing luciferase and ZsGreen (BEI catalog number NR-52516), a lentiviral helper plasmid expressing HIV Gag-Pol (BEI catalog number NR-52517), a lentiviral helper plasmid expressing HIV Tat (BEI catalog number NR-52518) and a lentiviral helper plasmid expressing HIV Rev (BEI catalog number NR-52519) along with a plasmid expressing spike protein of SARS-CoV-2 ^7^ using jetPRIME transfection reagent (Polyplus-transfection; NY, USA) as per manufacturer’s instruction. The culture supernatant containing pseudovirus particles was harvested at 48 hrs post-transfection, by centrifugation at 1200 rpm for 10 minutes and filtration through a 0.45 μM filter to remove cellular debris and then stored at −80° C freezer in aliquots for downstream applications.

The viral titers were determined using engineered HEK293T cells expressing the ACE2 receptor. For this purpose, 12,500 HEK293T-ACE2 cells were seeded per well in a poly-l-lysine-coated 96-well plate. 24 hrs after seeding, lentiviral particles were serially diluted with complete DMEM supplemented with polybrene (5mg/mL), and 50 μl of each dilution were added in four replicate wells. 48 hrs post addition, pseudoviral transduction efficiency was determined by measuring the activity of firefly luciferase in cell lysates using a bright-glo luciferase assay system (Promega, Madison, WI, USA; Cat# E2610). The luminescence was measured using a SpectraMax i3x multi-mode plate reader (Molecular Devices, San Jose, California, USA), and relative lluminescence units (RLUs) were plotted against virus dilutions.

### SARS-CoV-2 entry inhibitor screening assay

For screening SARS-CoV-2 entry inhibitors, 24 hrs before starting the assay, 20,000 HEK293T-ACE2 cells were seeded per well in a poly-l-lysine-coated 96-well plate. On the day of the assay setup, different concentrations of the compounds were mixed with the lentiviral particles expressing SARS-CoV-2 spike protein (1.00X10^5^ RLU/well) and incubated at 37°C for 30 min followed by the addition of 50 μl lentiviral particle-compound complex in the cells supplemented with polybrene (5mg/mL). After 48 hrs, the activity of firefly luciferase was measured as described above.

### Production and titration of SARS-CoV-2 stocks

SARS-CoV-2 isolate USA-WI1/2020 (BEI; cat# NR-52384) was passaged in Vero-STA-1 knockout cells. The viral titer was determined using the plaque assay ^54^. In brief, Vero E6 cells were seeded in 6-well plates. After 24 hours, cells were washed with sterile 1X PBS. The viral stock was serially diluted and added to cells in duplicate with new media and the plates were incubated at 37°C for 1 hour with occasional shaking every 15 minutes. Then, 2 mL of 0.5% agarose in minimal essential media (MEM) containing 5% FBS and antibiotics was added per well. Plates were incubated at 37°C for 72 hours. Then the cells were fixed with 4% paraformaldehyde overnight, followed by removal of the overlay and then staining with 0.2% crystal violet to visualize plaque-forming units (PFU). All assays were performed in a BSL-3 laboratory setting.

### Assessment of antiviral activity of selected compounds

The antiviral potential of the compounds were screened as reported earlier ^55^. In brief, UNCN1T and Vero-STAT1 knockout cells were seeded in 96 well plates 24 hours before infection at 20,000 cells/well. Different concentration of MU-UNMC-1 and MU-UNMC-2 ranging between 0.001 μM and 100 μM were added to the cells 2 hours prior to the infection. The cells were infected with 0.1 MOI of SARS-CoV-2 using Opti-MEM I reduced serum medium (Thermo Fisher, Cat#31985062) and incubated for 1 hour at 37° C with 5% CO_2_. For positive control, cells were treated with the same DMSO volume equivalent to the volume of drugs added. Mock-infected cells received only Opti-MEM^®^ I reduced serum medium. At the end of incubation of the virus, inoculum was removed, cells were washed with 1X PBS 3 times, and fresh media was added supplemented with the same concentration of drugs was added. Culture supernatant was collected at 24 hrs, and 48 hrs post-infection. The SARS-CoV-2 viral load was quantified in the culture supernatant using RT-QPCR with primer probes targeting the E gene of SARS-CoV-2 using PrimeDirect Probe RT-qPCR Mix (TaKaRa Bio USA, Inc) and Applied Biosystems QuantStudio3 real-time PCR system (Applied Biosystems, Waltham, MA, USA) per manufacturer’s instructions. Primers and probes used for SARS-CoV-2 RNA quantification were as follows: E_Sarbeco_F1: 5’ – ACAGGTACGTTAATAGTTAATAGCGT – 3’ (400nM), E_Sarbeco_R2: 5’ – ATATTGCAGCAGTACGCACACA – 3’ (400nM) and E_Sarbeco_P1: 5’ – FAM - ACACTAGCCATCCTTACTGCGCTTCG - BHQ1 – 3’ (200nM) as recommended by WHO. The SARS-CoV-2 genome equivalent copies were calculated using quantitative PCR (qPCR) control RNA from heat-inactivated SARS-CoV-2, isolate USA-WA1/2020 (BEI; cat# NR-52347). The percentage inhibition of SARS-CoV-2 replication in MU-UNMC-1 and MU-UNMC-2 treated cells was calculated with respect to viral loads in untreated control wells that received DMSO (considered 0% inhibition) and negative control wells (uninfected cells). IC_50_ values were calculated using four parameter variable slope sigmoidal dose-response models using Graph Pad Prism 8.0 software.

### Measuring the combinational antiviral potential of MU-UNMC-2 and Remdesivir

To determine the possible synergistic antiviral effect of MU-UNMC-2 on RDV and *vice versa* against SARS-CoV-2 replication, we tested combined doses of the two in SARS-CoV-2 infected UNCN1T and Vero-STAT1 knockout cells. The percentage inhibition of viral replication for each dose-combination was determined by RT-qPCR as described above. In brief, the UNCN1T and Vero-STAT1 knockout cells were seeded in 96-well plates at a density of 20,000 cells/wells 24 hrs before infection. Two hrs before infection, the cells were treated with different dose-combinations of MU-UNMC-2 and RDV, and then incubation with the drugs, cells were infected with 0.1 MOI of SARS-CoV-2. 24 hours post-infection, the supernatant was collected, and SARS-CoV-2 viral load was quantified using RT-qPCR as described above. The percentage inhibition of SARS-CoV-2 replication in the culture supernatant by different combination-doses was calculated with respect to viral load in positive control wells treated only with the DMSO (considered 0% inhibition) and negative control wells (uninfected cells). The percent inhibition of viral replication for 1:1 fixed-dose combination of the drugs was used to generate CI-Fa, isobologram, and dose-response plots. The combination index (CI) was calculated using multiple drug effect equations developed by Chou and Talalay using the CompuSyn algorithm (https://www.combosyn.com). CI values of <1 indicate synergy, values >1 indicate antagonism, and values equal to 1 indicate additive effects ^22, 56^. Dose-response percent inhibition matrix of single and combined treatment of RDV and MU-UNMC-2 in SARS-CoV-2 infected UNCN1T and Vero-STAT1 knockout cells 24 hrs post-infection and 3-D interaction landscape between Remdesivir and MU-UNMC-2 were calculated based on Loewe additive model using SynergyFinder v.2 ^57^.

### Statistical analysis

The CC_50_ and IC_50_ values were computed using four-parameter variable slope sigmoidal dose-response models using GraphPad Prism (version 8.0). The combination index (CI) was calculated using multiple drug effect equation developed by Chou and Talalay using the CompuSyn algorithm (https://www.combosyn.com). The 3-D interaction landscape between remdesivir and MU-UNMC-2 was calculated based on Loewe additive model using SynergyFinder v.2.

## Supplementary table legends

**Table S1: List of drugs like compounds selected for screening of in vitro antiviral activity against SARS-CoV-2.**

**Table S2: The median effective dose of MU-UNMC-2, remdesivir and their combination**

**Table S3: The CI values at a different simulated effective dose of fixed-dose combination of MU-UNMC-2 and remdesivir**

## Supplementary figure legends

**Supplementary Figure 1.**
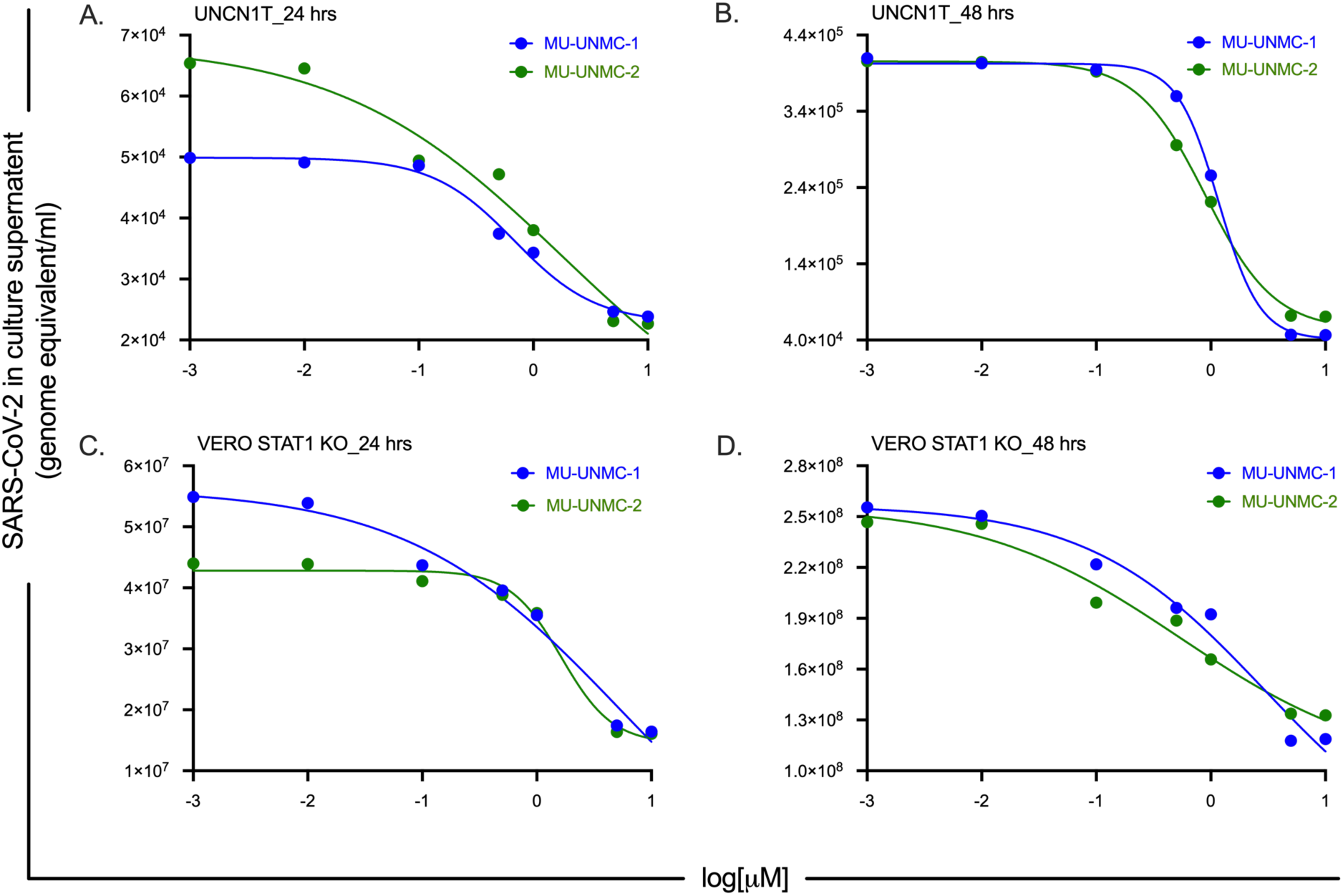
Replication kinetics of SARS-CoV-2 in MU-UNMC-1 and MU-UNMC-2 treated and SARS-CoV-2 infected UNCN1T cells and Vero-STAT1 knockout cells. (A & B) Real-time quantitative PCR data showing SARS-CoV-2 genome equivalent per ml of culture supernatant after 24 and 48 hrs post-infection in MU-UNMC-1 (in blue) and MU-UNMC-2 (in green) treated UNCN1T cells at indicated drug concentrations. (C & D) Real-time quantitative PCR data showing SARS-CoV-2 genome equivalent per ml of culture supernatant after 24 and 48 hrs post-infection in MU-UNMC-1 (in blue) and MU-UNMC-2 (in green) treated Vero-STAT1 knockout cells at indicated drug concentrations.

**Supplementary Figure 2:**
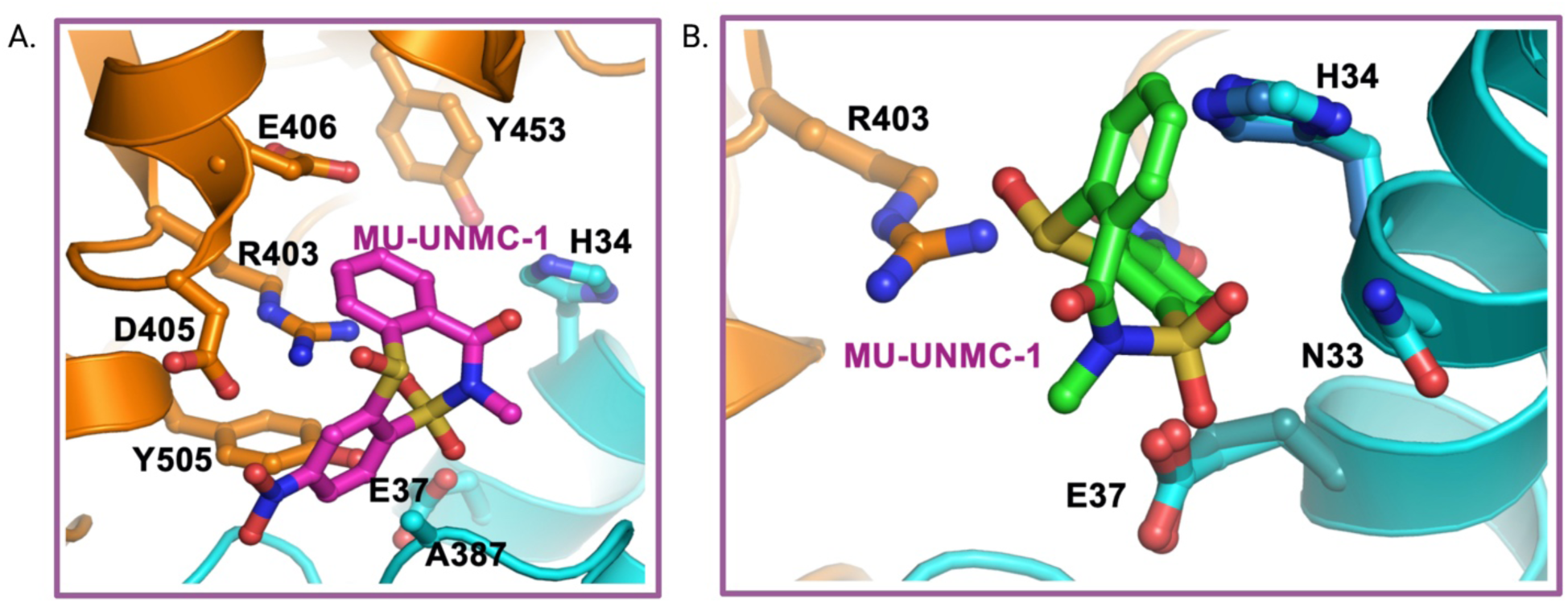
MU-UNMC-1 has two docking modes within the interface of SARS-CoV-2 spike and human ACE2 receptor.

**Supplementary Figure 3.**
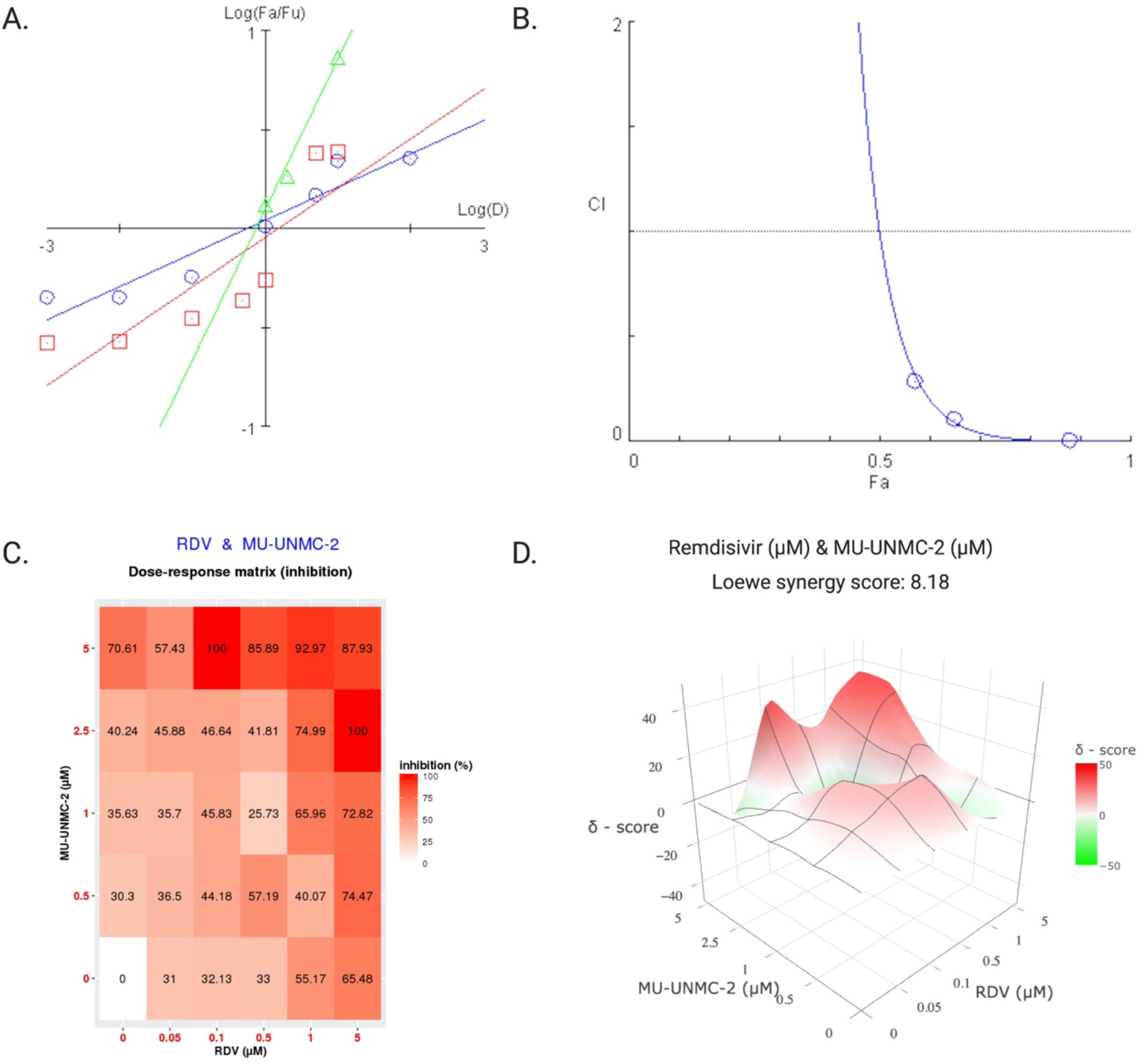
Synergistic effect of Remdesivir and MU-UNMC-2 combined treatment against SARS-CoV-2 in Vero-STAT1 knockout cells 24 hrs post-infection. (A) The Median-Effect plot (Chou plot) of MU-UNMC-2 (the red line with square data points), remdesivir (blue line with circular data points) and fixed dose combination of remdesivir and MU-UNMC-2 (green line with triangular data points). The median-Effect equation, that describes the dose-effect relationship is given by (Fa/Fu) = (D/Dm)^m^ or log(Fa/Fu) = m log(D) – m log(Dm). Fa is defined as percentage inhibition of viral growth at dose D and Fu is the fraction that remains unaffected (i.e., Fu = 1 - Fa). (B) Combination index plot or Chou-Talaly plot or Fa-CI plot: A plot of CI (combination index) on the Y-axis vs. Fa on X-axis for fixed-dose combination of Remdesivir and MU-UNMC-2 in SARS-CoV-2 infected Vero-STAT1 knockout cells (blue line with circular data points); (C) Dose-response percent inhibition matrix of single and combined treatment of Remdesivir and MU-UNMC-2 in SARS-CoV-2 infected Vero-STAT1 knockout cells 24 hrs post-infection. (D) 3-D interaction landscape between Remdesivir and MU-UNMC-2 calculated based on Loewe additive model using SynergyFinder v.2 in SARS-CoV-2 infected Vero-STAT1 knockout cells 24 hrs post-infection (Loewe synergy score 8.18; with most synergistic area score of 13.24).

