## supplementary tables for "Discovery and in-vitro evaluation of potent SARS-CoV-2 entry inhibitors"

**Table S1: List of drugs like compounds selected for screening of in vitro antiviral activity against SARS-CoV-2.**

| **Compound ID:** | **IUPAC Names:** | **Structure:** | **Glide score** |
| --- | --- | --- | --- |
| MU-UNMC-1 | 10-methyl-5-nitro-2λ⁴,9λ⁶-dithia-10-azatricyclo[10.4.0.0³,⁸]hexadeca-1(16),3(8),4,6,12,14-hexaene-2,9,9,11-tetrone | 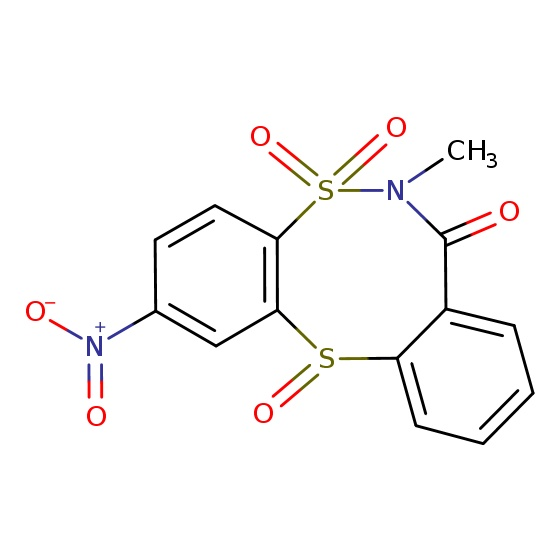 | - 9.8 |
| MU-UNMC-2 | 12-(trifluoromethyl)-2-oxa-6-thia-9-azatricyclo[8.4.0.0³,⁷]tetradeca-1(14),3(7),4,10,12-pentaen-8-one | 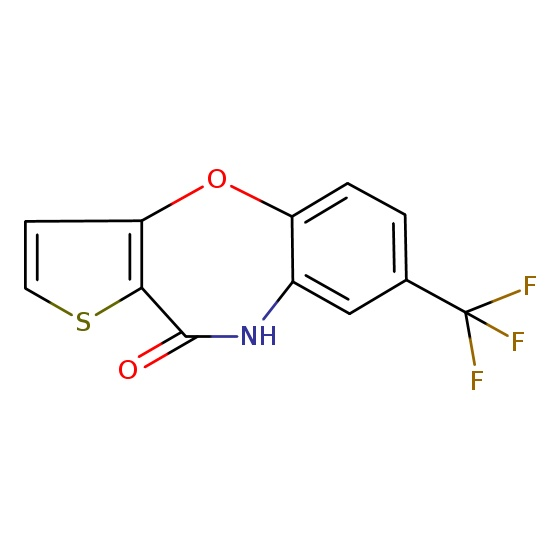 | - 11.6 |
| MU-UNMC-6 | 1-(3,4-dihydro-2H-quinolin-1-yl)-2-{[4-ethyl-5-(thiophen-2-yl)-1,2,4-triazol-3-yl]sulfanyl}ethanone | 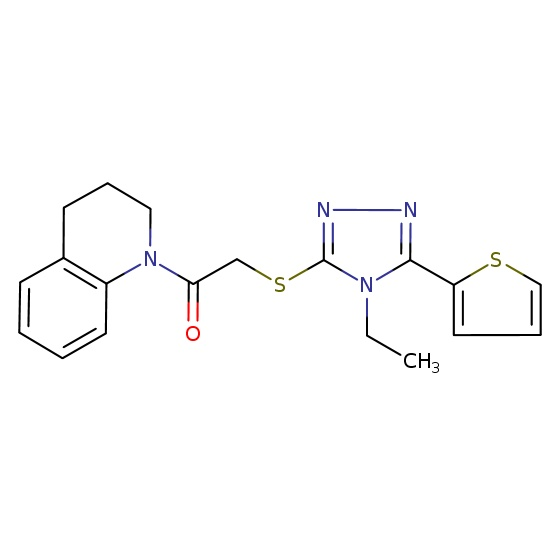 | - 7.6 |
| MU-UNMC-7 | 1-(indol-1-yl)-3-[(2-{[3-(trifluoromethyl)pyridin-2-yl]amino}ethyl)amino]propan-2-ol | 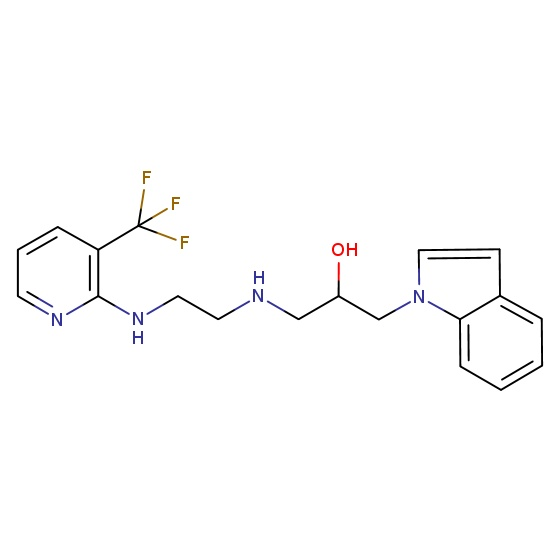 | - 7.2 |
| MU-UNMC-8 | (4Z)-2-[2-(4-chlorophenoxy)pyridin-3-yl]-4-[(dimethylamino)methylidene]-1,3-oxazol-5-one | 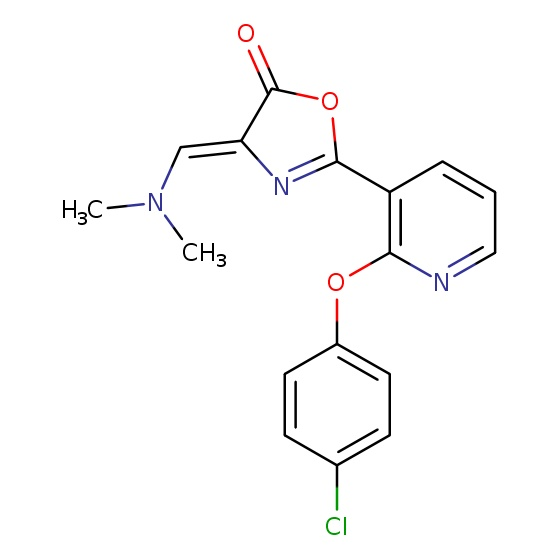 | - 6.4 |

**Table S2: The median effective dose of MU-UNMC-2, remdesivir and their combination**

| Vero-STAT1 knockout Cells | | | |
| --- | --- | --- | --- |
| Drug/Combo | Dm | m | r |
| MU-UNMC-2 (M-2) | 1.474 | 0.249 | 0.860 |
| Remdesivir (RDV) | 0.533 | 0.167 | 0.958 |
| Combination (RM-2) | 0.766 | 0.762 | 0.993 |
| UNCN1T Cells | | | |
| Drug/Combo | Dm | m | r |
| MU-UNMC-2 (M-2) | 3.454 | 0.369 | 0.957 |
| Remdesivir (RDV) | 0.272 | 0.231 | 0.949 |
| Combination (RM-2) | 0.244 | 0.786 | 0.968 |

Dm: The median effective dose; m: The slope of the median effect (ME) plot; r: The linear correlation coefficient of ME plot.

**Table S3: The CI values at a different simulated effective dose of fixed-dose combination of MU-UNMC-2 and remdesivir**

|  | CI values | | | |
| --- | --- | --- | --- | --- |
|  | ED_50_ | ED_75_ | ED_90_ | ED_95_ |
| Vero-STAT1 knockout cells | 0.9775 | 0.0176 | 7.14E-04 | 9.24E-05 |
| UNCN1T cells | 0.4830 | 0.0229 | 0.0020 | 5.72E-04 |
